# Structure of fungal tRNA ligase with RNA reveals conserved substrate binding principles

**DOI:** 10.1101/2024.06.06.597711

**Authors:** Sandra Köhler, Jürgen Kopp, Jirka Peschek

## Abstract

RNA ligases play a vital role in RNA processing and maturation including tRNA splicing, RNA repair and the unfolded protein response (UPR). In fungi and plants, the tripartite tRNA ligase Trl1 catalyzes the joining of TSEN-cleaved pre-tRNA exon halves. Trl1 also functions as ligase in the non-conventional *HAC1* mRNA splicing during the UPR. The final ligation step is performed by the N-terminal adenylyltransferase domain (LIG). The spatial arrangement of the exon ends during the ligation reaction has remained elusive. Here, we report the crystal structure of Trl1-LIG in complex with a tRNA-derived substrate. Our structure represents a snapshot of the activated RNA intermediate and defines the conserved substrate binding interface. The underlying enzyme-substrate interplay reveals a general substrate binding principle shared by adenylyltransferases. Moreover, we identify the determinants of RNA end specificity as well as the specific roles of Trl1-LIG’s subdomains during ligase activation, substrate binding and phosphoryltransfer.

## Main

The ligation of RNA ends by RNA ligases plays an important role during RNA maturation, processing and repair in all domains of life and viruses. Different classes of RNA ligases have evolved to fulfill a plethora of functions acting on specific subsets of RNA substrates^1^. Non-spliceosomal tRNA splicing is an essential step during tRNA biosynthesis^2^. In all eukaryotes, a set of pre-tRNAs contains an intervening intronic sequence located at one base 3′ from the anticodon^3^. Intron insertion at this canonical position disrupts the RNA tertiary structure of the anticodon stem-loop (ASL). This fully enzyme-catalyzed splicing reaction occurs in two steps: The intron is first excised by a tRNA-splicing endonuclease (SEN in yeast or TSEN in vertebrates), resulting in two exon halves that are subsequently connected by a tRNA ligase^4–6^. After endonucleolytic cleavage by the (T)SEN complex, the pre-tRNA exon halves contain a 2′,3′-cyclic phosphate (cP) at the 5′ exon and a 5′-hydroxyl (OH) group at the 3′ exon splice site^6–8^. Despite the conceptual similarity, the subsequent enzymatic reaction of tRNA exon-exon joining differs vastly depending on the organism^9^. In eukaryotes, there are two main branches: a fungal/plant as well as a metazoan/human strategy. The tRNA exon halves are ligated by the tRNA ligase Trl1 (previously named Rlg1) in fungi^5,10,11^ or RTCB as part of the multi-protein ligase complex in metazoans^8^. In addition, both ligases catalyze exon-exon ligation during the non-conventional splicing of the *HAC1/XBP1* mRNA as part of the unfolded protein response^12–15^. In fungi, the chemical modification at both RNA termini precedes the ligation of the pre-tRNA exon halves via formation of a new phosphodiester bond (Fig. 1A)^5^. The tripartite enzyme Trl1 catalyzes all ligation steps and consists of (1) an N-terminal adenylyltransferase domain (ligase; LIG), which belongs to the nucleotidyltransferase superfamily including DNA ligases and RNA capping enzymes, (2) a C-terminal cyclic phosphodiesterase domain (CPD) and (3) a central polynucleotide kinase (KIN) domain^11,16^. The CPD domain opens the 2′,3′-cP to form a 3′-OH/2′-phosphate (P) terminus, and the KIN activity phosphorylates the 5′-OH in a GTP-dependent reaction. Both modifications prepare the RNA substrate for the final adenylyltransferase reaction by the LIG domain. This ATP-dependent reaction progresses via three steps (Fig. 1A). First, the reaction between the active site lysine and ATP yields a covalent LIG-(lysyl-ζ)-AMP intermediate. Subsequently, the bound AMP is transferred to the 5′-P end of the 3′ exon and results via a pyrophosphate bond in a 5′-5′ RNA-adenylylate (AppRNA). Finally, the LIG domain catalyzes the nucleophilic attack by the 3′-OH on the RNA-adenylate to form a phosphodiester bond, thereby releasing AMP^5,11^. The remaining 2′-P at the splice junction is removed by the essential 2′-phosphodiesterase Tpt1^17–20^. The importance of a 2′-P for the adenylyltransferase reaction and its presence after ligation is a characteristic of Trl1-type RNA-ligases.

**Fig. 1:**
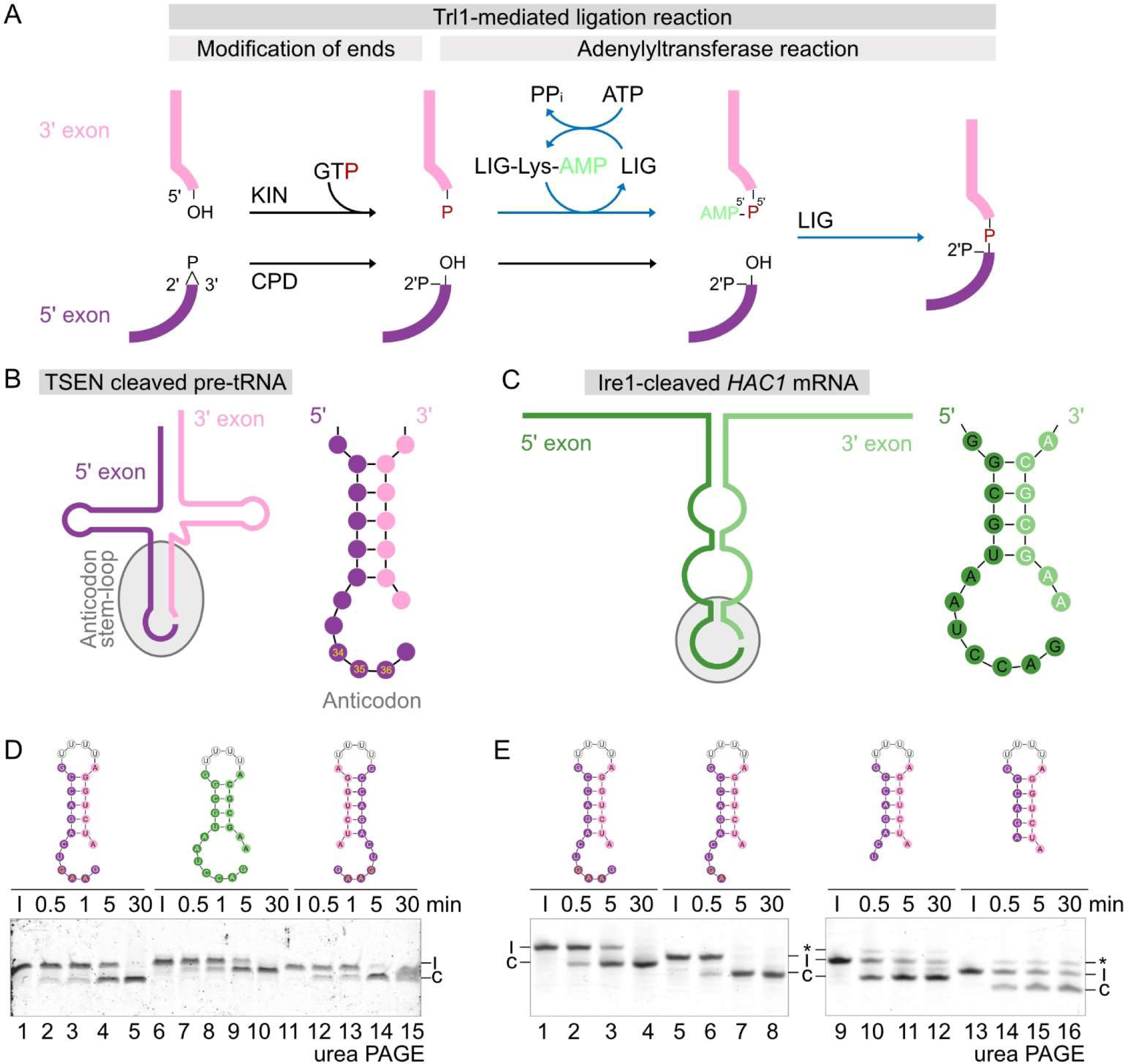
Trl1 ligation mechanism and RNA substrate structures. A, Overview of the tRNA ligation reaction by the fungal tRNA ligase Trl1. The RNA ends at the splice site of both exons (5′ exon in purple and 3′ exon in pink) are enlarged with the respective chemical groups at the termini. During the Trl1-catalyzed reaction the exon ends are first modified by the KIN and the CPD domain, followed by the adenylyltransferase reaction of the LIG domain resulting in phosphodiester bond formation. The same ligation mechanism joins the non-conventional *HAC1* mRNA exons during the unfolded protein response. B, The TSEN cleaved pre-tRNA halves are depicted as secondary structure with the 5′ exon (purple) and the 3′ exon (pink). The anticodon stem-loop (ASL) is highlighted with a grey boarder and enlarged on the right. The ASL folds in the same way for all cleaved pre-tRNAs: a 5 base-pair stem-loop with six unpaired nucleotides (including the anticodon triplet at position 34-36 in yellow) at the 5′ splice site and one unpaired nucleotide at the 3′ splice site. C, The IRE1 cleaved *HAC1* mRNA is depicted as secondary structure with the 5′ exon (dark green) and the 3′ exon (light green). The stem-loop with the splice site is highlighted with a grey boarder and enlarged on the right. The structure consists of a 4 base-pair stem-loop with seven unpaired nucleotides at the 5′ splice site and two unpaired nucleotides at the 3′ splice site. D, Circularization assay of *Ct*Trl1-LIG WT (1 µM) with the linear ASL-4U, HAC1-4U and ASL-4U^inv^ substrates. The ligation of the linear RNA substrate (l) to the circular form (c) was monitored by urea PAGE over time. E, Circularization assay of WT CtTrl1-LIG (0.1 µM) with varying length of the single-stranded overhangs at the 5′ exon 3′ end. Ligation of the linear ASL-4U substrates (l) to the circular form (c) as well as formation of the AppRNA intermediate (*) was monitored by urea PAGE.

Crystal structures of the homologous LIG domain from the thermophilic fungus *Chaetomium thermophilum (syn. Thermochaetoides thermophila*) have been solved^21,22^. The LIG domain has a bilobal architecture comprised of an N-terminal canonical adenylyltransferase domain (NTD) and a C-terminal domain (CTD) with a unique all-helical fold. This building principle of a conserved adenylyltransferase core with a unique C-terminal extension is also observed in other RNA and DNA ligases^23–25^. Previous structural and enzymatic data on the tRNA repair enzyme from Bacteriophage T4 Rnl1 show that its CTD is not required to catalyze RNA ligation in general but confers specificity for tRNA substrates^24^. In addition, the structure of the double stranded RNA editing ligase KREL1 from *Trypanosoma brucei*, likewise revealed a diverged CTD with an unknown function^25^. The specific role of the CTD within Trl1-type RNA ligases has remained elusive. Furthermore, reported structures of ATP-dependent RNA ligases revealed conformations associated with different nucleotide-bound states^24,26–29^. However, the existing structural information on nucleic acid binding and coordination of the exon ends is scarce. The crystal structure of the nick-sealing RNA ligase Rnl2 from the bacteriophage T4 bound to an adenylylated nicked DNA (App-DNA) duplex provides important insights into substrate binding by a dsRNA ligase^30^. In addition, DNA ligase structures with bound substrates (also with activated App-DNA) reveal binding modes of adenylyltransferases to nucleic acids^31–33^. However, the field is missing high-resolution structural insight into the coordination of physiological RNA substrates explaining the specificities of the individual RNA ligase families. Specifically, the coordination and orientation of both exon-ends in the case of RNA ligases acting on single-stranded RNA substrates remains elusive. Furthermore, the structural underpinnings of the specific requirement for a 2′-P of Trl1-type ligases are unknown. Here, we present the crystal structure of *Ct*Trl1-LIG in complex with an activated RNA substrate. Our structure reveals insights into substrate binding, activation and enzyme specificity. Complementing biochemical analyses dissect the role of the unique CTD and identify the determinants of 2′-P end specificity.

## Results

### Physiological Trl1 substrates consist of an RNA duplex stem with single-stranded overhangs

In order to understand the principles underlying LIG-RNA interaction and exon-exon coordination, we compiled the predicted RNA structures of both types of physiological substrates, TSEN-cleaved pre-tRNAs and Ire1-cleaved *HAC1* mRNA, respectively (Fig. 1A,B and Extended Data Fig. 1A).

Recent structures of the TSEN complex with pre-tRNA substrates indicate that cleaved pre-tRNAs assume the same overall fold as the mature tRNA^34–36^. There are six unpaired nucleotides (including the anticodon triplet) at the 5′ splice site and one unpaired nucleotide at the 3′ splice site (Fig. 1B). Since the position of the splice site is invariable in all eukaryotic cells^37^, all cleaved pre-tRNA substrates in *S. cerevisiae* consist of a 5-base-pair stem with single stranded overhangs (Extended Date Fig. 1A). In the case of *HAC1* mRNA, it was shown that base-pairing of complementary stretches keeps both exons together after cleavage by Ire1^38^. Similar to cleaved pre-tRNAs, in the predicted structure of *HAC1* mRNA, the ends at the splice site extend as overhangs from a duplex stem. There are seven unpaired nucleotides at the 5′ splice site and two unpaired nucleotides at the 3′ splice site (Fig. 1C).

In order to assess the minimal requirements of an RNA substrate and to determine the effect of RNA substrate structure on Trl1-LIG-catalyzed ligation, we tested a series of different RNA substrates derived from pre-tRNA anticodon stem-loops and the Ire1-clevead *HAC1* mRNA stem-loop (Fig. 1B and 1C). We designed and prepared a simplified tRNA anticodon stem-loop (ASL) substrate derived from yeast tRNA^Phe^ by linking the two exon strands with 4 uridines (4U) (Extended Data Fig. 1B; see Extended Data Table 2 for details). Likewise, a *HAC1*-stem-loop 4U-linked substrate (HAC1-4U) was designed. Ligation of either ASL-4U or HAC1-4U by *Ct*Trl1-LIG resulted in a circularized, faster migrating ligation product with similar kinetics (Fig. 1D, lanes 1-10). We noted that in both physiological substrates the single-stranded overhang at the 5′ splice site is longer in comparison to the 3′ splice site. To test, if the relative length of the single-stranded overhangs impacts ligation, we tested in vitro circularization with a pre-tRNA-derived substrate with inverted 5′/3′ overhangs (ASL-4U^inv^). *Ct*Trl1-LIG ligated ASL-4U^inv^ to completion with similar kinetics as the bona fide substrate (Fig. 1D, lanes 11-15).

We further tested if the length of the unpaired overhang at the 5′ splice site impacts ligation kinetics. To this end, we compared the canonical 6-nt overhang to substrates with 4-nt, 2-nt and 0-nt overhang (Fig. 1E). While the ASL-4U^-2nt^ RNA was ligated with faster kinetics, circularization of the ASL-4U^-4nt^ the ASL-4U^-6nt^ RNA was slower compared to the canonical 6-nt overhang. Moreover, we detected accumulation of the AppRNA intermediate for the two shorter overhangs (Fig. 1E, lanes 10-12 and 14-16, marked by *).

Additionally, we examined if LIG prefers a canonical anticodon stem-loop over an entirely single-stranded RNA (composed of A and G only) of the same length. Circularization (i.e. ligation) of the tRNA-derived ASL-4U RNA was faster compared to the unstructured RNA (Extended Data Fig. 2A, lanes 1-10). We noticed that the circularized ligation product of the structured RNA substrate migrated faster than the single-stranded A-G-only substrate of the same length despite the urea-denaturing conditions. Thus, we used an equivalent, folded stem-loop RNA with only A-U base-pairs in the stem area, which confirmed the underlying difference in migration behavior as well as the faster ligation kinetics for stem-loop substrates (Extended Data Fig. 2A, lanes 11-15). Using electrophoretic mobility shift assays (EMSAs) we revealed a slightly decreased affinity to the RNA without overhang (0-nt) compared to the canonical stem-loop substrate (Extended Data Fig. 2B).

Taken together, these biochemical analyses showed that a combination of RNA folding and overhang length at the splice sites determine the efficiency of exon-exon ligation.

### Structure of *Ct*Trl1-LIG with an activated RNA substrate

In order to gain structural insights into the binding and coordination of RNA substrates by the LIG domain, we aimed to crystalize *Ct*Trl1-LIG with an RNA substrate. Therefore, we screened various RNA substrates derived from the structure of both physiological Trl1 RNA substrates, cleaved pre-tRNA or *HAC1* mRNA. We focused our efforts on shortened RNA substrates to reduce the conformational flexibility outside of Trl1-LIG’s RNA binding cleft. Based on RNA secondary structure prediction^39^ and substrate requirement tests assessed during biochemical analyses (Fig. 1 and Extended Data Fig. 2), we specifically tested crystallization of *Ct*Trl1-LIG with various duplex-forming RNA oligonucleotides with short single-stranded overhangs in the presence of ATP. We obtained diffracting microcrystals belonging to space group P212121 for *Ct*Trl1-LIG (residues 1 to 414) with the first 7 nucleotides of the 3′ exon from cleaved yeast pre-tRNA^Phe^ with a 5′-P end (5′-P-AUCUGGA-3′). This RNA substrate was predicted to form a duplex due to self-complementarity with overhangs of two nucleotides at the 5′ end and one nucleotide at the 3′ end. We determined the LIG•RNA complex structure at 2.37 Å resolution using molecular replacement (Fig. 2A, see Table 1 for data collection and refinement statistics). Indeed, the single stranded RNA dimerized due to its five reverse complementary base pairs and crystallized as bound RNA duplex (Fig. 2A). Moreover, our structure contained bound AMP in the LIG active site in the same orientation as in previously solved structures. There was continuous electron density between the α-phosphate of AMP and the 5′-P of one of the RNA strands indicating the formation of an activated AppRNA ligation intermediate (Fig. 2B). The AMP moiety of the activated AppRNA intermediate is coordinated by residues T146, L147, K148 and K325. The continuing 3′ exon is further contacted by R99, K169 and K323. All of these residues are located within the NTD. While the ligase/RNA assembly within the unit cell promoted activation of the 5′ end, it did not position the 3′ end of the complementary strand correctly for ligation, and we did not observe any protein-RNA interactions to this strand (see Fig. 2A, sand-colored). We therefore attribute no physiological relevance to this strand, which rather served in stabilizing the activated strand to enable crystallization. We will describe additional physiologically relevant RNA binding modes in later sections.

**Fig. 2:**
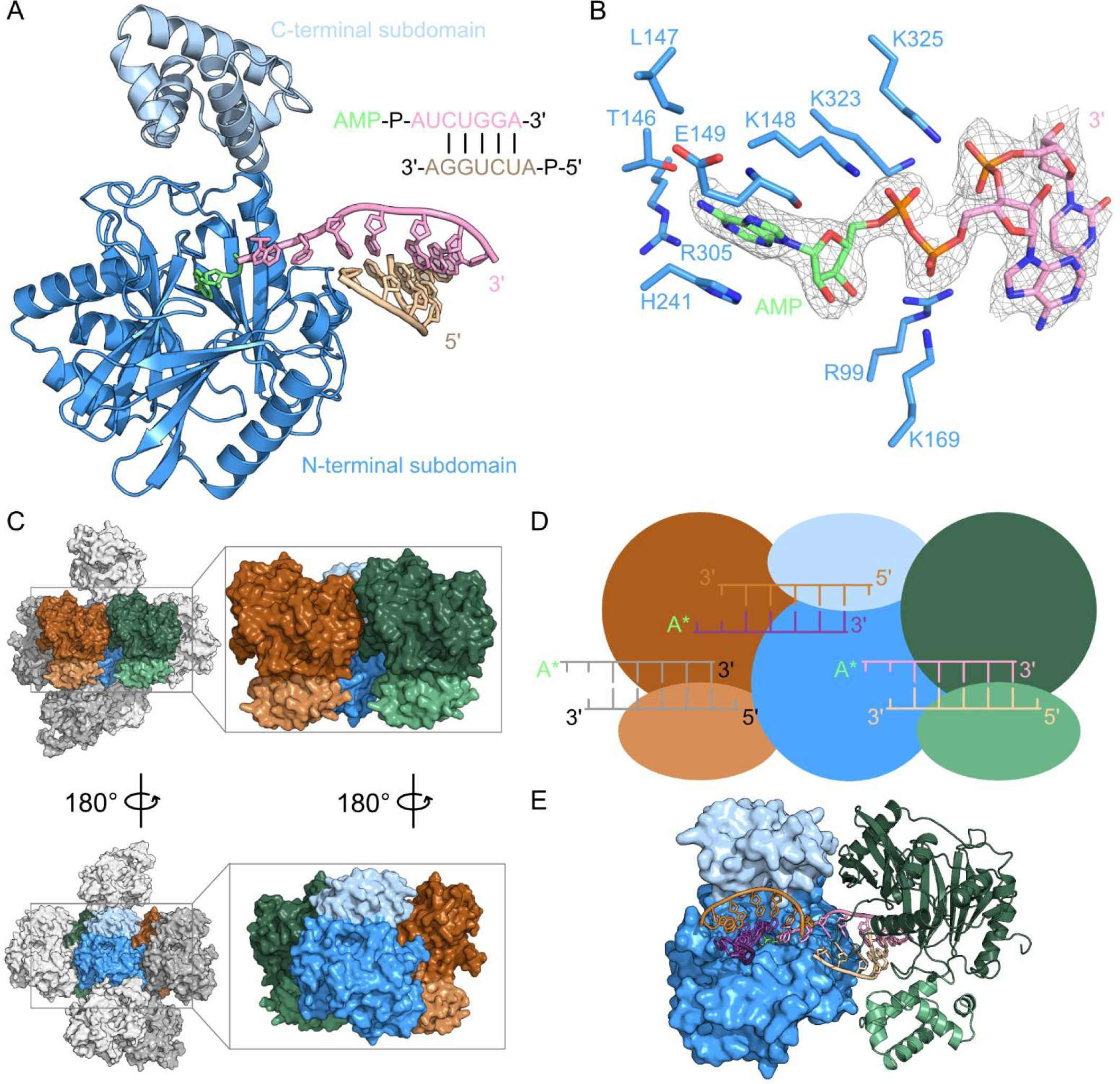
Structure of *Ct*Trl1-LIG with an activated RNA substrate. A, Cartoon representation of the overall structure of *Ct*Trl1-LIG in complex with an activated RNA substrate. The N-terminal domain is colored in blue and the C-terminal domain in light blue. The AMP cofactor (lime) is depicted as sticks and the RNA strands (pink and sand) in cartoon style. The RNA sequence overview contains the corresponding colors. B, Zoom into the active site with the activated AppRNA 5′ end and the coordinating amino acid residues depicted as sticks. The 2Fo-Fc electron density OMIT map at 1 σ is shown as a grey mesh. C, Nearest symmetry mates (relative to the blue molecule) of the LIG•RNA complex in the P212121 crystal. The individual molecules are shown in surface depiction. Two symmetry mates of the assembly (colored in orange and green) are enlarged for simplification (black rectangles). D, Scheme of the arrangement in C illustrating the end-to-end bridging of LIG molecules and their active sites by the bound RNA substrate molecules (same color code as in C). E, Surface representation of *Ct*Trl1-LIG (blue) with bound RNA duplexes (bronze and purple, pink and sand) as well as a neighboring symmetry mate in cartoon depiction (green) visualizing the binding of one RNA substrate between two LIG molecules in the crystal.

**Table 1.** Crystallographic data collection and refinement.

| CtTrl1-LIG•RNA |  |
| --- | --- |
| *PDB ID* | **8RBJ** |
| ***Data collection*** |  |
| Space group | P2<sub>1</sub>2<sub>1</sub>2<sub>1</sub> |
| Resolution (Å) | 2.37 - 47.6 (2.37 – 2.46) |
| Unique reflection | 20384 (2019) |
| Cell dimensions |  |
| a, b, c (Å) | 51.56, 75.36, 124.56 |
| α, β, γ (°) | 90, 90, 90 |
| R<sub>merge</sub> | 0.355 (3.123) |
| R<sub>pim</sub> | 0.090 (0.856) |
| Mean (I/σ(I)) | 7.7 (1.0) |
| Multiplicity | 15.8 (13.6) |
| Completeness (%) | 100 (100) |
| CC<sub>1/2</sub> | 0.989 (0.392) |
| ***Refinement*** |  |
| R<sub>work</sub> (%) | 17.87 (29.59) |
| R<sub>free</sub> (%) | 21.78 (37.04) |
| RMSD Bond length (Å) | 0.007 |
| RMSD Bond angle (°) | 0.950 |
| Ramachandran favored (%) | 97.4 |
| Ramachandran allowed (%) | 2.3 |
| Ramachandran outliers (%) | 0.3 |
| Rotamer outliers (%) | 1.2 |
| Clash score | 2.5 |
| Average B factor (Å<sup>2</sup>) | 60.3 |
| Protein | 60.4 |
| Ligands | 48.2 |
| Solvent | 47.7 |

Values in parentheses refer to the highest resolution shell.
The R<sub>free</sub> set consists of 5% randomly chosen data excluded from refinement.

In order to position the obtained LIG•RNA structure in the three-step ligation mechanism of the LIG domain, we compared with previously reported structures of *Ct*Trl1-LIG in the ATP/AMPcPP-bound as well as the activated LIG-Lys-AMP state (Extended Data Fig. 3A)^21,22^. Overall, the conformational changes in between the three states are subtle. Superposition with the adenylylated LIG (PDB ID: 6N0V) shows that the adenine base, ribose and αP of the AMP cofactor remain in place during adenylyl transfer (Extended Data Fig. 3B). The orientation of the broken and newly formed bonds at the αP support in-line attack. The phosphoryl transfer coincides with a stereochemical inversion at the αP center. The catalytic site lysine (K148 in *Ct*Trl1-LIG) shows minimal movement between both states. We conclude, that our LIG•RNA structure represents a state directly after adenylyl transfer from the active site lysine onto the 5′ RNA end (i.e. post step 2, see Extended Data Fig. 3B). The Mn^2+^ ion, which is present in the LIG-AMP structure, is absent in our LIG•RNA structure.

Noticeably, the co-crystallized RNA duplex did not show a simple 1:1-interaction with LIG. Instead, in the crystal lattice of the substrate complex, the RNA duplexes fill the space between neighboring LIG symmetry mates (Fig. 2C and 2D) by bridging two protein molecules in an end-to-end arrangement (Fig. 2E). This crystal packing allows for additional interpretations with regards to substrate binding from the viewpoint of one LIG molecule and all its RNA contacts within the crystal (Fig. 2E, blue LIG molecule). In the following, we describe and analyze the different RNA substrate binding modes in more detail.

### An extended RNA-binding cleft determines RNA end coordination

When we analyzed all RNA-protein interactions of a single LIG molecule within the crystal, we found that it binds two RNA duplexes in its extended substrate binding cleft (Fig. 3A). In addition to the aforementioned RNA duplex as defined in the ASU (Fig. 2A), a second RNA duplex occupies the opposite half of the substrate binding cleft (Fig. 3A,B). This binding mode suggests an additional dsRNA binding surface within LIG that utilizes both, the NTD and CTD to cradle the RNA duplex (Fig. 3A, duplex on the left). The CTD contacts one strand of the duplex with only one residue, N382 (Fig. 3C; compare to the bronze-colored RNA strand in Fig. 3A). The other strand of this additional duplex is coordinated by a cluster of residues within the NTD (Fig. 3A, purple strand). These NTD residues localize to an extended loop segment spanning residues 168 to 181. Parts of this loop form a thumb-like protrusion at one end of the substrate binding cleft of the NTD (Extended Data Fig. 3C). This loop region exhibits the highest conformational variability when comparing all available *Ct*Trl1-LIG structures (Extended Data Fig. 3A). Most RNA-binding residues contact the phosphate backbone (K64, Y68, R83, I153, S168, K169, H170, S171 and H182) while additional contacts to 2′-OH (R175 and H182) groups allow discrimination of RNA from DNA (Fig. 3C). We did not identify any interactions with the RNA bases.

**Fig. 3:**
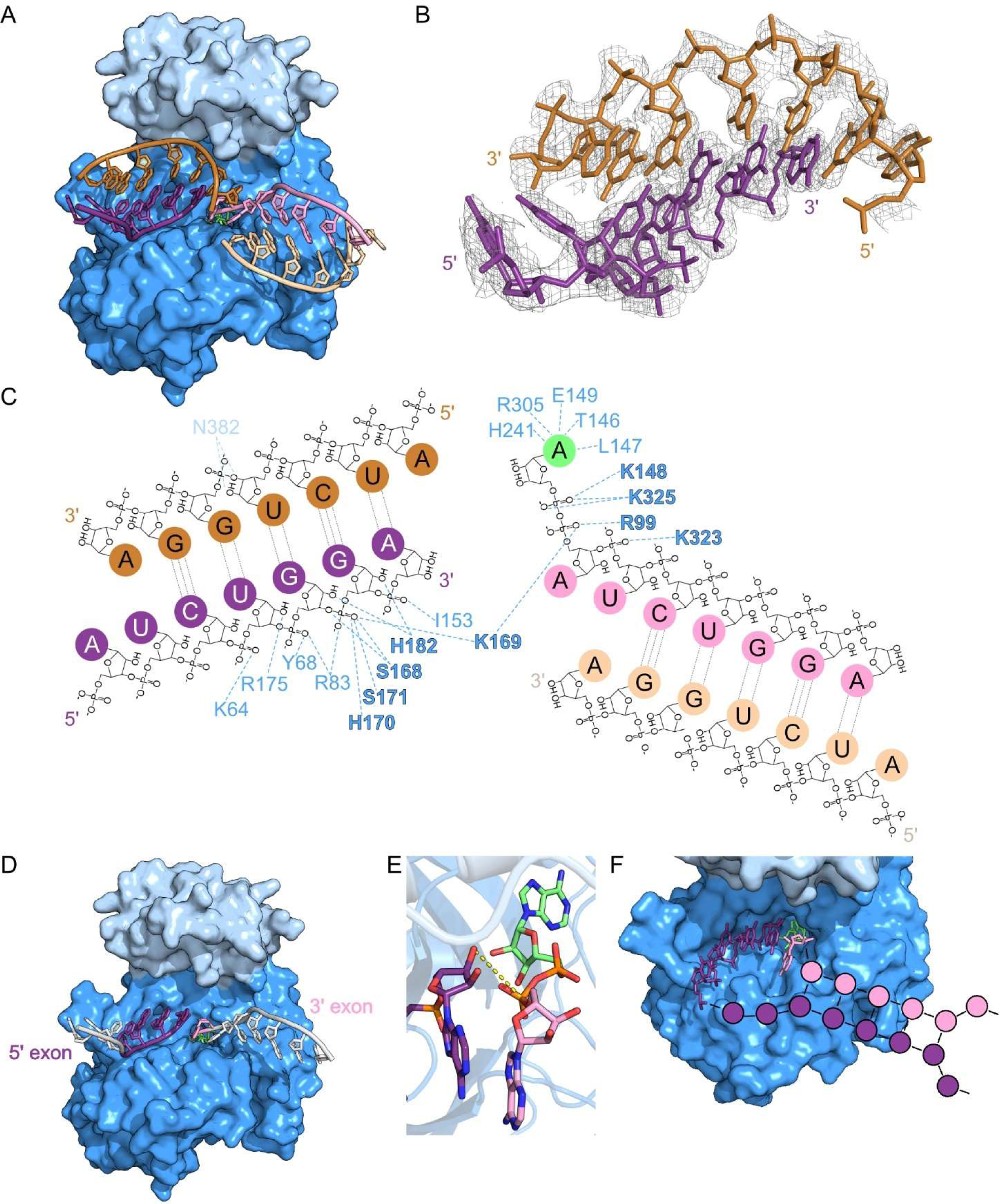
Analysis of the Trl1-LIG RNA binding interface. A, Surface representation of one *Ct*Trl1-LIG molecule with both interacting RNA duplexes in the crystal (see Fig. 2E). The N-terminal domain is colored in blue and the C-terminal domain in light blue. The RNA strands (bronze and purple; sand and pink) are depicted as cartoon with the AMP cofactor as sticks in lime. B, Zoom into the RNA duplex forming strands (bronze and purple) depicted as sticks. The 2Fo-Fc electron density OMIT map at 1 σ is shown as a grey mesh. C, Interaction map of *Ct*Trl1-LIG residues (one letter code) with the RNA duplexes from the NTD (blue) and the CTD (light blue) using the same color coding as in Fig. 2B and 3B. Essential amino acid residues are shown in bold. D, Representation of the exon-exon arrangement extracted from the structure of *Ct*Trl1-LIG•RNA. The RNA exon strands are annotated as 5′ exon (purple) and 3′ exon (pink). RNA nucleotides without amino acid contacts are shown in white. E, Zoom into the *Ct*Trl1-LIG active site with the 3′ end of the 5′ exon (purple, end colored by element) and the activated 5′ end of the 3′ exon (pink, end colored by element, AMP in lime) depicted as sticks. The distance of 4.7 Å between the 3′-OH and the 5′-P is indicated by the yellow dashed line. F, Scheme of pre-tRNA anticodon stem-loop coordination. Nucleotide positioning from the crystal structure are shown as sticks in the active site of *Ct*Trl1-LIG (blue). The extending nucleotides are depicted as dots building the anticodon stem-loop of a pre-tRNA (5′ exon in purple and 3′ exon in pink).

By focusing on the RNA ends that are oriented towards the active site to allow formation of a new phosphodiester bond (step 3 of the ligation reaction) as well as involved in LIG/RNA interactions, we interpret the ultimate 5′ adenylylated nucleotide (Fig. 3D, pink RNA nucleotide) as the 3′ exon end equivalent and, consequentially, the four nucleotides of the strand from the other duplex, which is coordinated by the NTD with its 3′ end reaching into the active site, as the 5′ exon equivalent (Fig. 3D, purple RNA strand). The remaining nucleotides of the two strands did not show any interactions with LIG (Fig. 3D, white RNA strands). The distance of 4.7 Å between the 3′-OH and the αP as well as their relative orientation suggest that additional re-positioning of the 3′ end is required for the final ligation step (Fig. 3E, zoom). This notion is in line with our interpretation of the LIG•RNA complex representing a post-step 2 state.

Alignment-based conservation analysis^40,41^ revealed that the entire substrate binding cleft of Trl1-LIG including all RNA-interacting residues within our LIG•RNA structure is evolutionary conserved (Extended Data Fig. 4). In order to determine the in vivo relevance of the residues involved in RNA binding, we used yeast plasmid-based mutagenesis. If these residues are essential for the enzyme activity, cells are not viable anymore. In a previous study, a mutagenesis screen tested viability of *Saccharomyces cerevisiae* (*S. cerevisiae, Sc*) cells for alanine substitutions of many residues that interact with the RNA in our structure^42^ For the remaining residues, we substituted the homologous amino acid residues in yeast to alanine and tested for growth (Extended Data Fig. 5). With the exception of I153, R175 and N382, the identified RNA-binding residues are essential for cell survival, reinforcing our findings in the context of physiological relevance (Fig. 3C, bold labels).

Based on our structural data and the composition of a cleaved pre-tRNA anticodon stem-loop with six unpaired nucleotides (including the anticodon triplet) at the 5′ splice site and one unpaired nucleotide at the 3′ splice site (Fig. 1B), we schematically depicted the orientation of the anticodon stem-loop. (Fig. 3F).

### The substrate-binding mode is conserved between Trl1-LIG and nick-sealing adenylyltransferases

Up to now, the structure of T4 Rnl2 with a nicked nucleic acid substrate represents the only high-resolution snapshot of an RNA ligase engaged with its substrate^30^. T4 Rnl2 is a nick-sealing ligase whose function is the repair of single-strand breaks in the context of double-stranded RNA segments. In this structure, the bound substrate is a DNA duplex with a terminal 2′-OH and an adenylylated 5′ DNA end at the nick. It revealed two distinct forms of the complex representing (1) the state immediately after 5′ end adenylylation (A form, step 2 product) as well as (2) the state before nucleophilic attack by the 3′-OH (B form, step 3 substrate). We employed structural superposition based on the core adenylyltransferase domain of our LIG•RNA complex with the T4 Rnl2-substrate complex to compare the substrate binding modes and specific RNA end positioning. When superimposing both structures by the β-sheets within the core adenylyltransferase domain, the three DNA strands in the Rnl2 structure, i.e. both nicked strands and the complementary template strand, have equivalents in our LIG•RNA structure (Fig. 4A). Importantly, the nick 5′ end is adenylylated and superimposes to the AMP-3′ exon strand in our structural model. Both ligases utilize the same set of conserved lysine and arginine residues to coordinate the adenylylated 5′ end (Fig. 4B). When comparing our structure to both complex forms of Rnl2 with bound DNA, the strand harboring the 3′ nick strand as well as the complementary template strand superimpose with one bound RNA duplex in our structure. (Fig. 4C). The adenine base and ribose of the co-factor nucleotide superimpose with both complex forms of Rnl2. The position of the αP as well as the orientation of the phosphoanhydride bond are more similar to the post step 2 state (complex A) in line with our aforementioned interpretation. Moreover, superposition with DNA ligase structures of the human DNA ligase 1^32^ and the *E. coli* LigA protein^31^ with adenylylated nicked substrate revealed a similar overall architecture of ligase/substrate complexes (Extended Data Fig. 6). Taken together, comparison of both RNA ligase/substrate complexes confirmed the existence of two RNA binding modes within Trl1-LIG: (1) an RNA duplex-binding interface as well as (2) the specific coordination of both RNA exon ends towards the catalytic center.

**Fig. 4:**
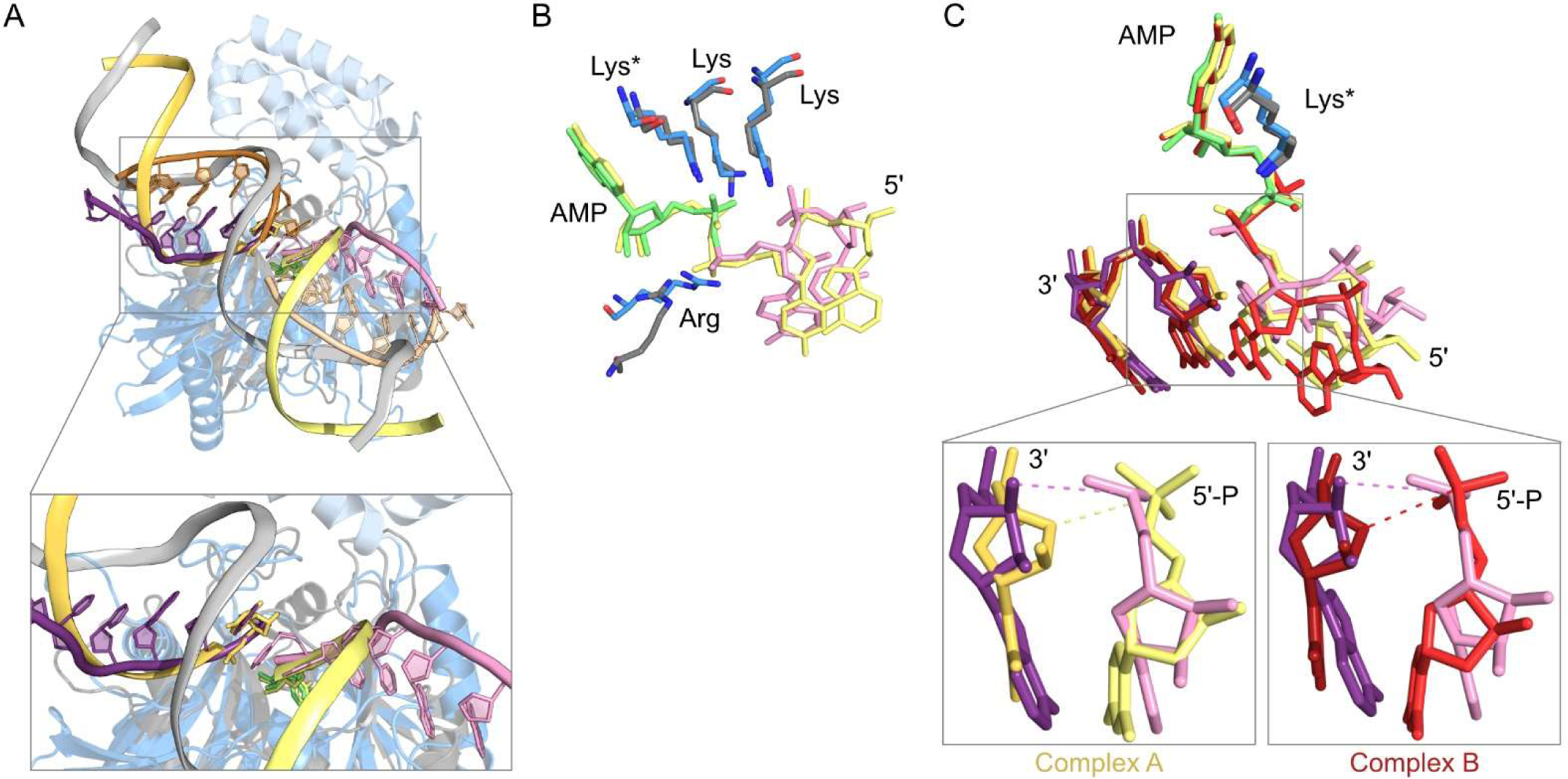
Conserved RNA binding modes between *Ct*Trl1-LIG•RNA and T4 Rnl2•AppDNA. A, Superposition of the *Ct*Trl1-LIG•RNA structure and the structure of T4 Rnl2 (grey) in complex with an AppDNA substrate (yellow, template strand in grey; PDB ID: 2HVR, A form). The structures were superimposed by the core β-sheets of the adenylyltransferase domain. The DNA strands are depicted as cartoon strands. The rectangles show the enlarged view. Note the omission of one RNA strand (compare Fig. 3A, bronze strand). B, Zoom into the activated 5′ end of the nucleic acids after superposition of the structures from A depicted as sticks. The active site lysins (Lys*, K148 of *Ct*Trl1-LIG and K35 of T4 Rnl2) as well as the conserved coordinating lysins (Lys K323 and K325 of *Ct*Trl1-LIG, K225 and K227 of T4 Rnl2) and arginines (Arg, R99 of *Ct*Trl1-LIG and R55 of T4 Rnl2) are shown as sticks. C, Zoom into the activated 5′ end of the nucleic acids and the 3′ end of the corresponding 5′ exon after superposition of the structures from A as well as an alternative conformation of the T4 Rnl2 complex with an AppDNA substrate (red; PDB ID: 2HVR, B form). The active site lysins are depicted as sticks (K148 of *Ct*Trl1-LIG and K35 of T4 Rnl2) together with the first two nucleotides of both exon strands. The individual superpositions are depicted on the bottom (rectangles). Dashed lines indicate the distance of the 3′ end to the 5′-P (pink: 4.7 Å, yellow: 4.4 Å, red: 3.6 Å).

### Role of the C-terminal subdomain during Trl1-LIG-mediated ligation

The CTD of Trl1-LIG adopts a unique fold unlike any other adenylyltransferase^21,22^ and was shown to be essential for ligase activity in vivo^43^. However, our substrate complex structure identified only limited contacts of residues from the CTD to the bound RNA, which motivated us to identify its role during exon-exon ligation. To this end, we generated a shortened version of *Ct*Trl1-LIG with only the NTD (ΔCTD; aa 1-328) and determined its effect on each catalytic step of the Trl1-LIG-catalyzed ligation reaction (Fig. 1A). First, we analyzed the contribution of the CTD during the required adenylylation of Trl1-LIG (i.e. activation step). To this end, we monitored the formation of the covalent LIG-K148-AMP intermediate using radioactive [αP^32^]-ATP with WT LIG as well as the K148N and ΔCTD variants (Fig. 5A). WT LIG showed concentration-dependent adenylylation indicated by the occurrence of a specific band in the autoradiogram, which was completely absent for the K148N variant. The degree of adenylylation for the ΔCTD variant was indistinguishable from the WT protein, which we confirmed in a time-course assay (Extended Data Fig. 7). Our data reveal that the CTD is dispensable during ligase activation.

**Fig. 5:**
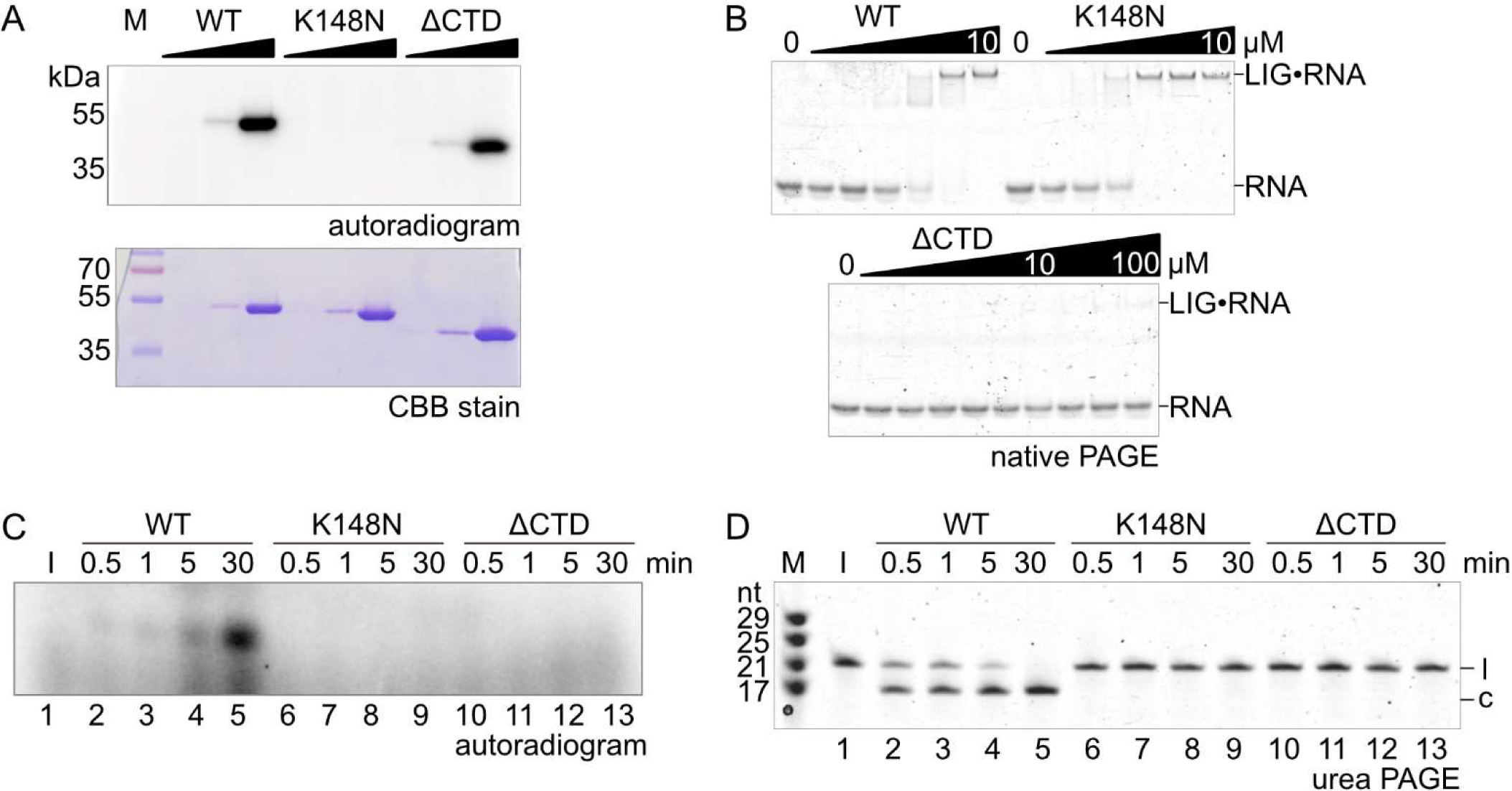
Role of the Trl1-LIG C-terminal domain during RNA ligation. A, Adenylyltransferase assay of *Ct*Trl1-LIG WT, K148N and ΔCTD variants. The upper panel shows the autoradiogram after incubation of increasing protein concentrations (0.1-10 µM) with αP^32^-labeled ATP. The SDS-PAGE was Coomassie Brilliant Blue (CBB) stained (lower panel) as protein loading control. B, Electrophoretic mobility shift assay (EMSA) with *Ct*Trl1-LIG WT, K148N (both 0-10 µM) and ΔCTD variants (0-100 µM). Increasing amounts of protein were incubated with the ASL-4U RNA (1 µM) and analysed by native PAGE using SYBRGold RNA staining. Formation of the LIG•RNA complex was assessed by band shift. C, RNA adenylylation by *Ct*Trl1-LIG WT, K148N and ΔCTD variants (1 µM). The autoradiogram shows formation of the adenylylated ASL-4U intermediate via αP^32^-AMP incorporation over time. D, Circularization assay of *Ct*Trl1-LIG WT, K148N and ΔCTD variants (1 µM). Ligation of the linear ASL-4U substrate (l) to the circular form (c) was monitored by urea PAGE.

The second step of the Trl1-LIG-mediated ligation reaction is binding of the RNA exons. We determined the substrate binding capacity of the different LIG variants using EMSAs with the ASL-4U substrate (Extended Data Fig. 1B). Deletion of the CTD strongly reduced RNA binding compared to WT and K148N, which both showed stable complexes at similar concentrations (Fig. 5B). With ΔCTD, we only observed detectable levels of binding at a 100-fold higher enzyme concentration compared to WT. The EMSA revealed that the CTD contributes to RNA binding.

In the next step of the ligation reaction the LIG-bound AMP forms a covalent anhydride with the 5′-P end of the RNA resulting in an AppRNA intermediate. To monitor the formation of this adenylylated RNA intermediate, we performed ligation reactions with the ASL-4U RNA in the presence of [αP^32^]-ATP. The autoradiogram revealed signals for radioactively labeled RNA in the presence of WT LIG (Fig. 5C, lanes 2-5). No activated RNA intermediate was detected, when the K148N (Fig. 5C, lanes 6-9) and ΔCTD variants (Fig. 5C, lanes 10-13) were used. These adenylylation reactions were monitored over 30 min. With increasing reaction time, the signal for the AppRNA intermediate using WT LIG subsided due to completion of the ligation reaction and concomitant release of AMP (Extended Data Fig. 8A,B). When using the ASL-4U substrate with non-ligateable 3′ end (2′-OH/3′-P, 5′-P), we observed continuous increase in adenylylation over the same time-course (Extended Data Fig. 8C,D).

Lastly, we tested the final ligation step in circularization assays. In the presence of WT LIG, the linear substrate was completely circularized within 30 min (Fig. 5D, lanes 2-5). Addition of ΔCTD or the ligase-dead K148N variant revealed no ligation/circularization activity under the same conditions (Fig. 5D, lanes 6-13). This series of in vitro biochemical assays revealed that the CTD was dispensable during ATP-dependent activation of Trl1-LIG. However, presence of the CTD promoted RNA binding and was required for transfer of the AMP moiety from the active-site lysine onto the 5′ RNA end as well as the final ligation step. Taken together, our in vitro biochemical assays rationalize the in vivo importance of the CTD.

### The CTD provides 2′ phosphate specificity via conserved arginine residues

The requirement for a terminal 2′ phosphate is a specific characteristic of Trl1-type RNA ligases^5,44^. The 2′ phosphate is not present in our crystal structure of LIG•RNA, but we can infer an approximate location of the 2′-P from the terminal 2′-OH end and its position near the active site (Fig. 6A). Interestingly, this position is occupied in previous *Ct*Trl1-LIG structures by a sulfate ion^21,22^. Superposition of the LIG•RNA complex structure and the LIG•AMPcPP structure (PDB ID: 6N67) depicts the potential 2′-P pocket (Fig. 6B). The sulfate ion is coordinated by two conserved arginines within the CTD, R334 and R337 (see conservation scale in Extended Data Fig. 4), which are both located in proximity of the RNA 3′ end in our LIG•RNA structure (Fig. 6A). To assess the role of R334 and R337 in 2′-P specificity, we purified *Ct*Trl1-LIG variants R334A, R337A as well as the double mutant R334A/R337A and determined their activity towards different 2′/3′ RNA ends. To this end, we performed ligation assays based on circularization of the ASL-4U substrate harboring a 5′-P in combination with a 2′-P/3′-OH (the same as endogenous substrates), a 2′-OH/3′-OH or a 2′-OH/3′-P end. The assays revealed slower ligation kinetics for all three Arg-to-Ala variants towards the endogenous-like RNA in comparison to WT LIG. Reduced ligation kinetics of the variants coincided with an accumulation of the AppRNA intermediate (Fig. 6C, upper panel). Both observed effects were stronger in the R334A variant compared to R337A and were additive in the double mutant. The same assay with the 2′-OH/3′-OH,5′-P ASL-4U substrate (note the non-physiological 2′-OH end) resulted in overall diminished circularization activities with a slightly increased activity of the R334A variant compared to WT, R337A and R334A/R337A (Fig. 6C, middle panel). The correct identity of the substrate ends in these assays was confirmed by using T4 Rnl1, which did not ligate the 2′-P/3′-OH-containing ASL-4U but only its canonical 2′-OH/3′-OH-containing substrate. In a control experiment, using the ASL-4U with 2′-OH/3′-P,5′-P ends none of the ligases showed circularization (Fig. 6C, bottom panel). We observed formation of the AppRNA intermediate for 3′ exon fragment even when combined with non-productive 5′ exon fragments albeit with reduced kinetics (Fig. 6C, middle and bottom panel, marked with *). Taken together, these results indicate a strong preference but not a strict requirement for 2′-P-containing substrates by *Ct*Trl1-LIG, which is strongly dependent on the CTD residue R334.

**Fig. 6:**
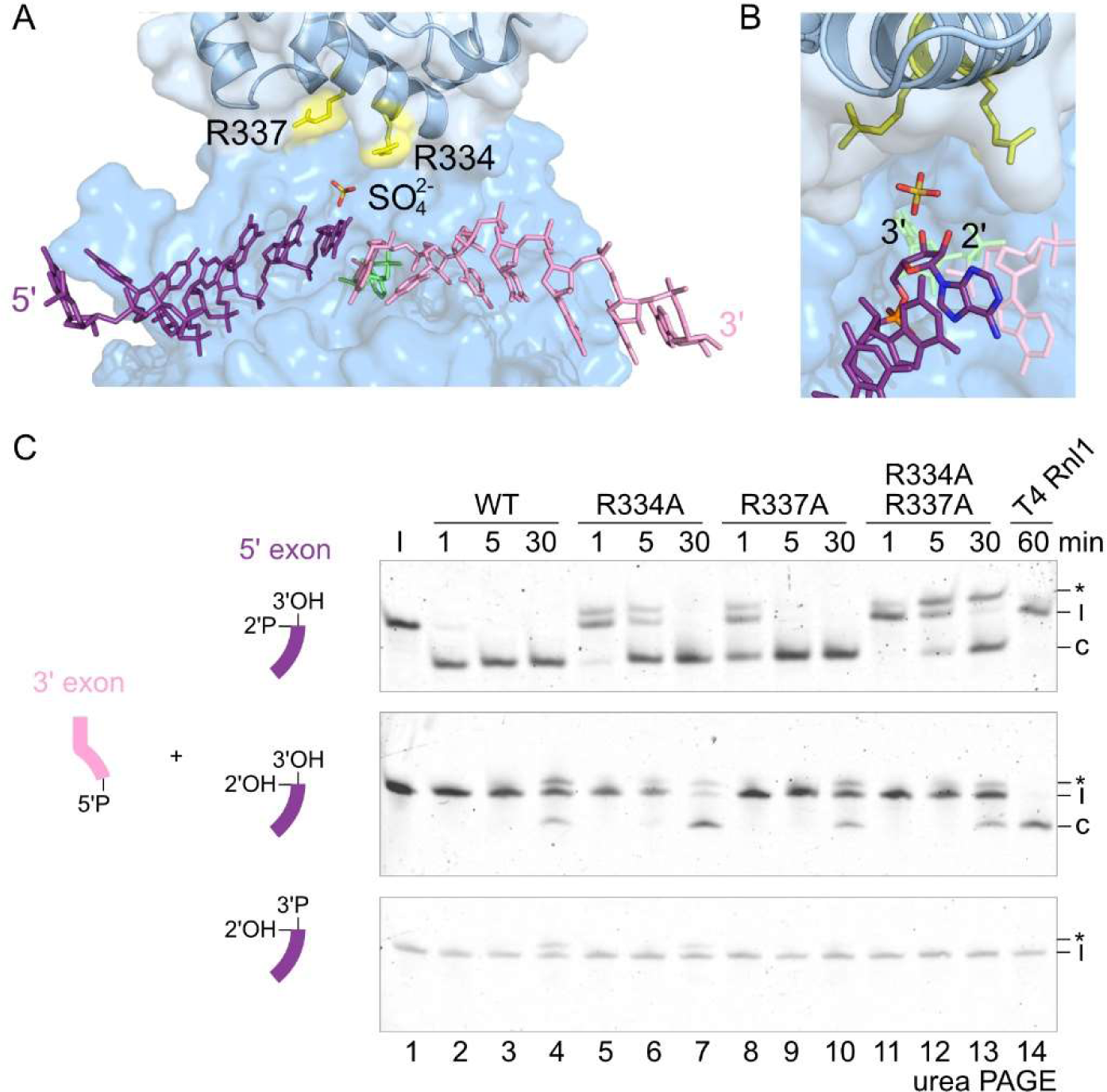
Conserved arginine residues within the CTD mediate 2**′-**P specificity. A, The position of a bound sulfate ion (orange) from the previous crystal structure of *Ct*Trl1-LIG in complex with AMPcPP (PDB: 6N67) is shown in the context of the *Ct*Trl1-LIG•RNA structure. Depiction of the sulfate ion based on structural alignment of both crystal structures. The CTD residues R334 and R337 (yellow) were previously found to coordinate the sulfate ion. The RNA strands (purple and pink) with the bound AMP (lime) are depicted as sticks. B, Zoom into the 3′ RNA end coordination. The sulfate ion is located in a potential 2′-P pocket in close proximity to the 2′-OH end of the 5′ exon end (purple, colored by element). C, Circularization assay of *Ct*Trl1-LIG WT, R334A, R337A and the R334A-R337A double mutant (1 µM) with the ASL-4U substrate and the indicated end modification on the 2′ and 3′ ends of the 5′ exon. T4 Rnl1 was used as ligation control (single timepoint). Ligation of the linear ASL-4U substrates (l) to the circular form (c) with the AppRNA intermediate (*) was monitored by urea PAGE.

## Discussion

The fungal tRNA ligase Trl1 is an essential enzyme for both tRNA splicing and non-conventional *HAC1* mRNA splicing during the unfolded protein response^5,12^. Despite recent structural progress, the principles of RNA substrate recognition by Trl1 as well as the coordination of the exon ends by its ligase domain remain unknown. Overall, our understanding of RNA ligases that join single-stranded RNA substrates is limited by the scarce availability of ligase-RNA structures. Here, we present the crystal structure of *Ct*Trl1-LIG with an RNA substrate. Our data provide high-resolution insights into coordination of the RNA exons by Trl1-LIG including a conserved RNA-binding surface for double-stranded RNA segments and identify molecular determinants for the specific biochemical mechanism at the exon ends.

By comparing our LIG•RNA structure to previously reported structures of both RNA and DNA ligases with an activated nucleic acid substrate, we identified a conserved binding mode for ATP-dependent activation of the 3′ fragment (or exon) also present in ssRNA ligases. We surmise that orientation of the activated strand relative to the core adenylyltransferase domain is mostly invariable between adenylyltransferases. Relative to the activated strand, the 5′ fragment (exon) is coordinated in apical orientation so that the entire RNA binding cleft is being utilized. With regards to the coordination of both RNA strands, the comparison with the substrate complex of the nick-sealing RNA ligase Rnl2 is particularly informative as it reveals a shared overall binding mode. Thereby, we uncovered a common RNA substrate recognition and binding principle shared between nick-sealing adenylyltransferases (both RNA and DNA ligases) as well as RNA ligases that join single-stranded breaks.

In contrast to other RNA ligase families, the requirement (or strong preference) for a 2′-P is a defining characteristic of Trl1-type ligases^5,44^. Recent structural studies suggested two arginine residues within the CTD of fungal Trl1-LIG (R334 and R337 in *Ct*Trl1, R295 and R298 in *Sc*Trl1), which coordinate a sulfate ion in place of a phosphate, as determinants for the 2′-P end specificity^21,22^. The LIG•RNA structure proposes an approximate position for the 2′-P based on the bound 5′ exon equivalent, which is indeed in the vicinity of the CTD arginines. Our biochemical assays confirm the proposed function of both CTD arginine residues. We conclude that the Arg-to-Ala variants show an accumulation of the AppRNA intermediate due to perturbed positioning of the 3′-OH via the 2′-P interaction with the arginines. While the impact of the R334A variant was stronger compared to R337A in our in vitro experiments, yeast mutagenesis of the equivalent R295 and R298 to alanine revealed only R337 to be essential in vivo^42^. Conceivably, both arginines act together in 2′-P binding, thereby orienting the 3′-OH end in a position favorable for nucleophilic attack of the cognate 5′ end. However, the precise role of each arginine remains to be elucidated. Noticeably, both arginine residues are conserved within fungi but absent in plant Trl1 ligases. In contrast to *Arabidopsis thaliana* Trl1, which requires the presence of a 2′-P for adenylylation of the 5′-end (i.e. step 2 of the LIG mechanism), fungal Trl1-LIG does not require a 2′-P for activation according to our data^44^. Both, the AppRNA formation during crystallization as well as our biochemical analyses corroborate the deviating 2′-P requirement between plant and fungal Trl1-LIG. Given the lack of plant Trl1-LIG structures, we speculate that their CTD fold, and thus the mode of 2′-P coordination, differs from the fungal counterparts. The presence of varying C-terminal extensions or subdomains is a common feature of adenylyltransferase-type RNA ligases. These C-terminal segments are not part of the core ATP-dependent ligase domain, but rather function in promoting distinct specificity dependent on the cellular substrates^23–25^. We show that the CTD of Trl1-LIG is not required for ligase activation (i.e. adenylylation), but plays a crucial role for RNA exon-exon ligation by promoting RNA binding. Since the aforementioned 2′-P-binding arginines are located in the CTD domain of Trl1-LIG, we conclude that its importance lies in the correct positioning of the 5′ exon RNA strand.

None of the RNA-binding residues contact specifically any RNA bases in the LIG•RNA structure. Instead, they coordinate the strands on the phosphate backbone, explaining why this enzyme is able to ligate a variety of RNA substrates independently of their sequence identities as long as they possess the correct ends, namely a 2′-P/3′-OH and a 5′-P. This ability is reflected in the diversity of cellular substrates (all intron-containing tRNAs and *HAC1* mRNA) as well as virtually unrestricted activity towards in vitro substrates. In addition, specific contacts with 2′-OH groups indicate the enzyme’s preference for RNA over DNA.

Circularization assays with different RNA substrates suggest that Trl1-LIG favors the exon ends held in close proximity by double stranded segments – as in an anticodon stem-loop. We surmise that base pairing of the two exon ends prior to ligation ensures the joining of the correct exon ends from the same RNA. Both canonical in vivo substrates, SEN cleaved pre-tRNAs as well as Ire1-cleaved *HAC1* mRNA exhibit such exon-exon base-pairing^34–36,38^. This RNA-intrinsic feature ensures joining of the correct exon ends despite the potential presence of other RNA fragments in the cell and favors ligation of the cleaved substrates over degradation by exonucleases^22,45^. Our structural insights into RNA coordination do not show binding of an entirely physiological substrate (e.g. cleaved pre-tRNA). Nevertheless, they provide insights into the coordination of both exon termini and identify important RNA-binding residues. Taken together, the LIG•RNA structure expands the collection of snapshots representing the individual steps of Trl1-LIG reaction mechanism.

The mechanism of substrate transfer within the full-length, tripartite Trl1 remains unknown. How are both exon-ends passed to the LIG domain after being modified by the KIN and CPD domain? We do not know if the exon ends are simultaneously or sequentially modified. Structural insights into the coordination of the RNA substrate by the KIN and CPD domains are currently missing, but would promote our understanding of the interplay between all three Trl1 active sites during catalysis.

Finally, Trl1 is often discussed as potential target for antifungal therapy due to the fundamental difference compared to the human tRNA ligase complex. In this study, we provide additional insights on how to potentially perturb RNA binding and also harness 2′-P specificity. In summary, our study uncovers conserved RNA binding modes between nick-sealing and single strand-end joining RNA ligases, indicating how different RNA ligase families evolved from a common principle to perform their specific cellular tasks.

## Methods

### Site directed mutagenesis

The pET15b-*Ct*Trl1-LIG plasmid^22^ served as template for PCR-based site direct mutagenesis with one oligonucleotide for K148N and E328-stop (ΔCTD) variants or two oligonucleotides and Q5^®^ Site-Directed Mutagenesis Kit (NEB) to introduce R334A, R337A mutations and R334A/R337A double mutation. Oligonucleotides used for mutagenesis are listed in Extended Data Table 1. Positive constructs were transformed in *E. coli* BL21-CodonPlus (DE3)-RIPL (Agilent Technologies) and used for protein expression as described below.

### Expression and purification of recombinant *Ct*Trl1-LIG WT and variants

In this study, recombinant His6-tagged *Ct*Trl1-LIG (residues 1-414) and variants were expressed in *E. coli* BL21-CodonPlus (DE3)-RIPL (Agilent Technologies). Cells were grown in Luria broth medium including ampicillin at 37°C until an OD600 0.8, expression was induced by 1 mM isopropyl-1-thio-b-D-galactopyranoside (IPTG), cells were grown for 3 h at 30°C and harvested by centrifugation. The cells were lysed in lysis buffer [25 mM HEPES/NaOH pH 7.5, 500 mM NaCl, 1 mM DTT, 25 mM imidazole, 1x cOmplete^TM^ protease inhibitor cocktail (Roche), and 10 mM Na4P2O7 for de-adenylylated state purification] using a microfluidizer. Purification was performed by Ni^2+^ affinity chromatography using a HisTrap FF column (GE Healthcare Life Sciences) in elution buffer (25 mM HEPES/NaOH pH 7.5, 500 mM NaCl, 1 mM DTT, 500 mM imidazole). After Tag removal in cleavage buffer (25 mM HEPES/NaOH pH 8.0, 150 mM NaCl, 2.5 mM CaCl2) by thrombin protease (Cytiva), the protease was removed from (un)cleaved target protein using a HisTrap FF column (GE Healthcare Life Sciences). Remaining nucleic acids were removed using anion exchange chromatography (AIEX) on a HiTrap Q HP 5 mL column with a NaCl gradient in high salt buffer (25 mM HEPES/NaOH pH 8.0, 1 M NaCl, 1 mM DTT). Further, purification proceeded by size-exclusion chromatography (SEC) using a HiLoad 16/60 Superdex200 pg column (GE Healthcare Life Sciences) in SEC buffer (10 mM HEPES/NaOH pH 7.5, 150 mM NaCl, 0.5 mM TCEP). Proteins were concentrated using Amicon Ultra-15 Centrifugal Filter Units 30 kDa MWCO (Merck).

His6-tagged *Ct*Trl1-CPD (residues 576–846) and His6-tagged *E. coli* RtcA were expressed and purified as described previously^22^.

### Crystallization

For crystallization of His6-*Ct*Trl1-LIG in complex with an RNA oligonucleotide (5′-P-AUCUGGA-3′), the protein (final concentration of 5.8 mg/mL) and RNA were mixed in a 1:1.7 (protein:RNA) ratio and used for sitting drop vapor diffusion at 291 K. Microcrystals appeared after 10 days in 0.2 M potassium formate and 20% PEG 3350. Microcrystals were flash-frozen in liquid nitrogen using 20% ethylene glycol as cryoprotectant.

### Data collection, structure determination and analyses

For His6-*Ct*Trl1-LIG-WT•RNA duplex, data sets were collected at cryogenic temperature at ESRF beamline ID23-2 from multiple small crystals in one single loop using the *MeshAndCollect* workflow^46^. Within the *MeshAndCollect* workflow, diffraction images of partial data sets were integrated with XDS^47^, clustered and combined, and finally scaled using AIMLESS^48^ as part of the CCP4i software package^49^. The resolution cut-offs were selected based on the half-data set correlation CC1/2 as implemented in AIMLESS^50^. Phases were obtained by molecular replacement with PHASER-MR^51^ implemented in the PHENIX package^52^. A previously solved structure of *Ct*Trl1-LIG (residues 1-414; PDB ID 6N67) served as search model. Iterative model building and refinement was performed with Coot^53^ and Phenix.refine^54^. The quality of the resulting structural models was analyzed with MolProbity^55^. Structure Figures were prepared with PyMOL 2.4.1 (The PyMOL Molecular Graphics System, Schrödinger, LLC.). Crystallographic data are summarized in Table 1. Coordinates and structure factors are deposited at the Protein Data Bank PDB with accession code 8RBJ.

### Radioactive adenylylation assay

Adenylylation reaction was performed with 0.1, 1 or 10 µM protein, 10 nM ATP and 18.5 MBq/mL [alpha]P^32^-ATP in assay buffer (20 mM HEPES/NaOH pH 7.5, 70 mM NaCl, 2 mM MgCl2, 1 mM TCEP, 5% glycerol) at 30 °C. The reaction was stopped at different timepoints in 4x loading dye (1M Tris, pH 6.8, 350 mM SDS, 50% glycerol, 3.7 mM bromophenol blue, 25% β-mercaptoethanol), separated on a 12% SDS-PAGE, fixed in fixation solution (20% ethanol, 7% acetic acid), incubated overnight on a radioactivity detecting film and scanned using a FLA 7000 (FUJIFILM).

### Electrophoretic mobility shift assay

Electrophoretic mobility shift assays (EMSAs) were used to detect protein-RNA interaction complexes. 1 µM of RNA was mixed with increasing concentrations of protein (0.04-10 µM or 25-100 µM) and 100-fold excess of ATP. Samples were incubated for 10 min at 30°C and stopped in native loading dye (0.5x TBE, 5 mM EDTA, 5% glycerol, trace amounts of xylene cyanol and bromophenol blue). Samples were loaded on a 10% non-denaturing PAGE and run in 0.5x-TBE buffer for 45 min at 100 V and 4°C. The gels were subsequently stained with SYBR Gold nucleic acid stain (Invitrogen) to visualize the RNA-containing complexes by ultraviolet trans-illumination.

### Radioactive RNA adenylylation detection

To detect adenylylated RNA intermediate circularization assay was performed in presence of radioactively labeled ATP. The yeast tRNA^Phe^ stem-loop derived RNA substrate ASL-4U (Extended Data Table 2) was diluted in RNA buffer (20 mM HEPES/NaOH pH 7.5, 100 mM NaCl and 1 mM MgCl2), thermally unfolded at 90°C for 1 min and refolded at room temperature by cooling down to∼40°C. Prior circularization, the 3′ end of the RNA was end-modified by incubating 2 µM of RNA with 0.5 µM RtcA, 0.5 µM *Ct*Trl1-CPD, 10 nM ATP and 18.5 MBq/mL [alpha]P^32^-ATP in RNA assay buffer (20 mM HEPES/NaOH pH 7.5, 70 mM NaCl, 2 mM MgCl2, 1 mM TCEP, 5% glycerol) for 30 min at 30°C. Addition of 1 µM *Ct*Trl1-LIG or -variants initialized the reaction, resulting in adenylylation of the RNA for the circularization of the stem-loop RNA over time. The reaction was quenched in 10-fold excess of stop solution (10 M urea, 0.1% SDS, 1 mM EDTA, trace amounts of xylene cyanol and bromophenol blue). RNA samples were unfolded at 80°C for 2 min. The samples were separated on a 15% denaturing urea PAGE. The gel was incubated overnight on a radioactivity detecting film and scanned using a FLA 7000 (FUJIFILM).

### *In vitro* circularization assay

The ASL-4U RNA substrate and variants as well as the HAC1-4U substrate (Extended Data Table 2) were used for circularization to detect ligation activity of *Ct*Trl1-LIG-WT and variants. The RNA was diluted in RNA buffer (20 mM HEPES/NaOH pH 7.5, 100 mM NaCl and 1 mM MgCl2), thermally unfolded at 90°C for 1 min and refolded at room temperature by cooling down to ∼40°C. Prior circularization, the 3′ end of the RNA was end-modified by incubating 2 µM of RNA with 0.5 µM RtcA, 0.5 µM *Ct*Trl1-CPD and 0.01 mM ATP in RNA assay buffer (20 mM HEPES/NaOH pH 7.5, 70 mM NaCl, 2 mM MgCl2, 1 mM TCEP, 5% glycerol) for 30 min at 30°C. To ensure correct folding after the modification reaction, the truncated RNA substrates were refolded again. Addition of 1 µM *Ct*Trl1-LIG or variants initialized the ligation reaction, resulting in circularization of the stem-loop RNA over time. The reaction was quenched in 10x excess of stop solution (10 M urea, 0.1% SDS, 1 mM EDTA, trace amounts of xylene cyanol and bromophenol blue). RNA samples were unfolded at 80°C for 2 min and subsequently placed on ice to prevent refolding. The samples were separated on a 15% or 12.5% denaturing urea PAGE and visualized via SYBR Gold nucleic acid stain (Invitrogen).

### Yeast strains and growth conditions

All yeast strains in this study were based on *Saccharomyces cerevisiae* W303 that was made *TRP1* and *ADE2* by repairing the endogenous auxotrophy and are listed in Extended Data Table 6. Yeast transformations were performed by the standard LiOAc/single stranded carrier DNA/PEG method, cells were recovered on YPD medium and grown on the appropriate selection medium. To test the impact of several Trl1-LIG mutation on yeast cell survival, the *trl1*Δ strain rescued by plasmid-based, constitutive (ADH promoter) expression of *Ct*Trl1^22^ was used for plasmid shuffle with different pRS313-*Sc*Trl1-3xHA constructs. The shuffle vector was cloned using Gibson assembly (NEBuilder Kit) after PCR amplification (Extended Data Table 3) of three DNA fragments with complementary overhangs: *Sc*Trl1-3xHA (from a synthetic gene block) as well as 1kb upstream and downstream of the *TRL1* gene (from genomic DNA). The fragments were cloned into the pRS313 vector^56^ using restriction site cloning with EcoRI and SmaI. The 3xHA tag was introduced after the C-terminal domain of Trl1 for co-immunoprecipitation experiments not shown in this study. Mutations within the pRS313-*Sc*Trl1-3xHA were introduced by site-directed mutagenesis (Extended Data Table 4) and confirmed by sequencing. The *trl1*Δ strains rescued by the p416ADH-*Ct*Trl1 plasmid were transformed with the respective shuffle vectors (Extended Data Table 5). Counterselection on 5-fluoroorotic acid (FOA) containing SDC plates was used to test for growth of the yeast strains with the introduced mutations in the *TRL1* gene.

For growth tests under different selection conditions, five-fold serial dilutions were spotted onto either standard SDC-Ura, SDC-His or SDC/FOA plates. Respective images are shown in the respective Figure (Extended Data Fig. 5).

### Denaturing polyacrylamide gel electrophoresis

Circularization reactions were carried out as described above, and the reaction products were loaded onto a 12.5% or 15% urea gel. The gels were run in 1x TBE buffer at 150 V (constant voltage) at room temperature for 70 min. The gels were subsequently stained with SYBR Gold nucleic acid stain (Invitrogen) to visualize the RNA by ultraviolet trans-illumination.

### Native polyacrylamide gel electrophoresis

EMSA samples were prepared as described above and loaded on a 10% native PAGE. The gels were run in cold 0.5x TBE buffer at 100 V (constant voltage) at 4°C for 45 min. The gels were subsequently stained with SYBR Gold nucleic acid stain (Invitrogen) to visualize the RNA by ultraviolet trans-illumination.

### Sodium dodecyl sulfate polyacrylamide gel electrophoresis

Adenylylation reactions were carried out as described above. The reaction products were loaded onto a 12% SDS-PAGE or a 12% NuPAGE (Invitrogen) using 4x loading dye (1 M TRIS, pH 6.8, 350 mM SDS, 50% glycerol, 3.7 mM bromophenol blue, 25% β-mercaptoethanol). The gels were run in 1x SDS-Running-buffer (248 mm TRIS base, 1.92 M glycine, 1% SDS) at 250 V (constant voltage) at room temperature for 35 min or in 1x MOPS-SDS buffer (50 mM MOPS, 50 mM TRIS, 0.1% SDS, pH 7.7). The gels were subsequently stained with Coomassie to visualize the proteins and de-stained with 10% acetic acid solution.

## Acknowledgments

We are grateful to Michael Brunner for fruitful discussion. We acknowledge the European Synchrotron Radiation Facility (ESRF) for provision of synchrotron radiation facilities and we would like to thank Romain Talon and Max Nanoa for assistance in using beamline ID23-2 and data processing. We acknowledge support from the BZH Crystallization Platform and thank Claudia Siegmann for help with protein crystallization. We thank Klemens Wild for valuable discussions. The authors gratefully acknowledge the data storage service SDS@hd supported by the Ministry of Science, Research and the Arts Baden-Württemberg (MWK) and the German Research Foundation (DFG) through grant INST 35/1314-1 FUGG and INST 35/1503-1 FUGG. JP acknowledges funding from the Deutsche Forschungsgemeinschaft (DFG, German Research Foundation) – Emmy Noether Program (project number 442512666) and TRR319 TP-A02 (project number 439669440).

## Author contributions

S.K. performed experiments and analyzed data. J.K. and J.P. analyzed crystallographic data. J.P. conceived the study and planned the experiments. The manuscript was written by S.K., J.K. and J.P.

