## Extended data for "Structure of fungal tRNA ligase with RNA reveals conserved substrate binding principles"

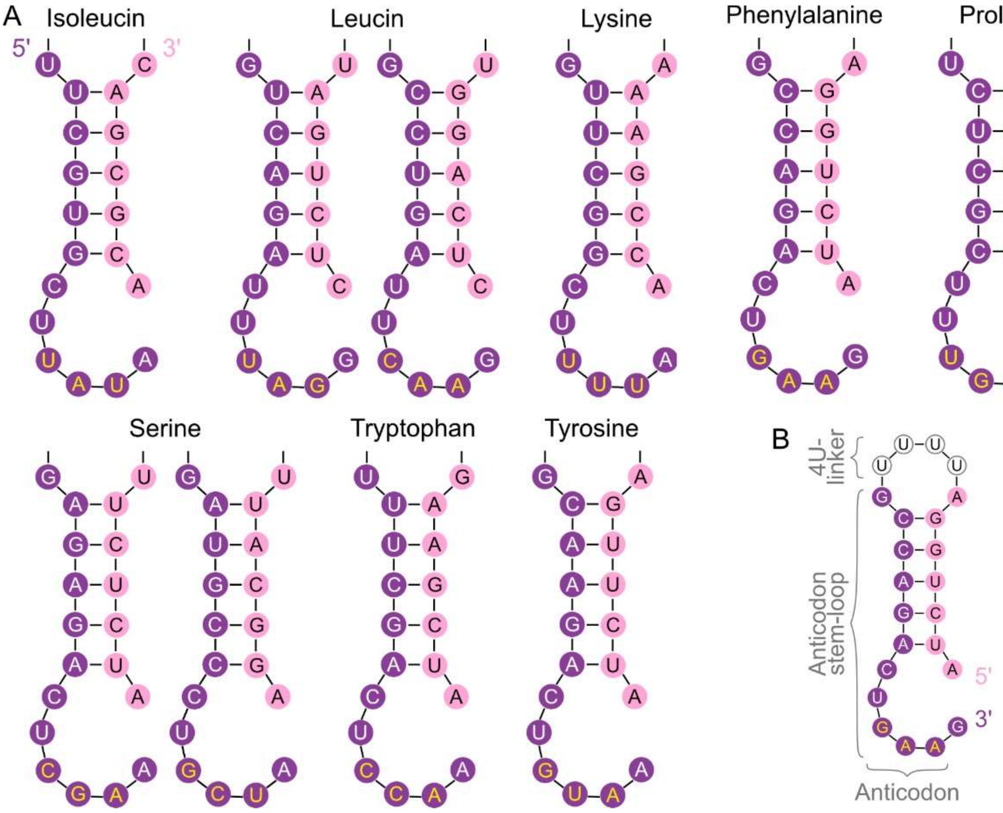


**Extended Data Fig. 1: Structures of TSEN cleaved pre-tRNAs and circularization assay substrate.**

A, Yeast tRNA anticodon stem-loop structures. Overview of the anticodon stem-loop regions of all intron-containing tRNAs in yeast. The 5′ exon is colored in purple and the 3′ exon in pink. The sequences for the individual tRNA anticodon stem loops are depicted with the corresponding anticodon in yellow letters.

B, Scheme of the ASL-4U RNA substrate for circularization, adenylylation and binding assays. The RNA is derived from the TSEN-cleaved yeast $tRNA^{Phe}$ anticodon stem-loop (ASL) with both strands linked by four uridines (4U).

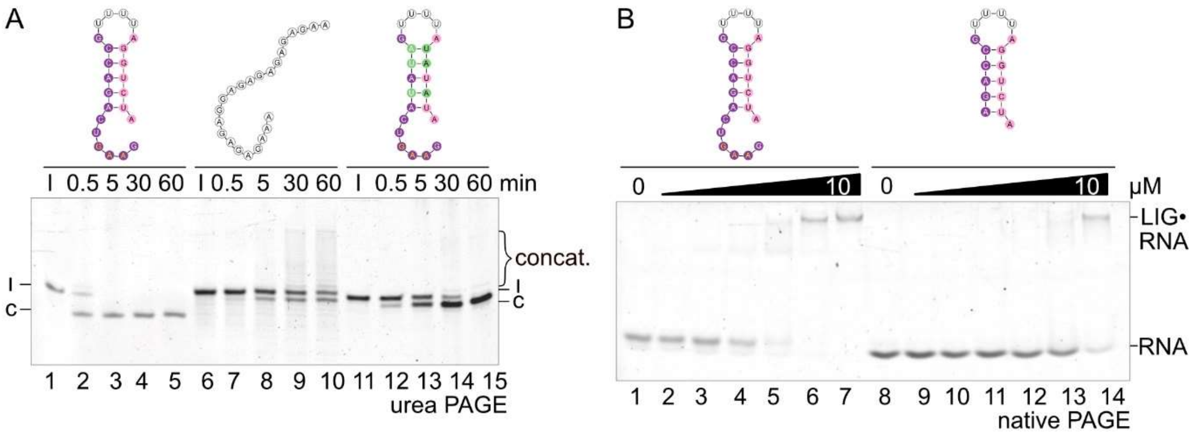


**Extended Data Fig. 2: RNA stem-loop structure determines *Ct*Trl1-LIG activity.**

A, Circularization assay of WT *Ct*Trl1-LIG (1 µM) with structured and unstructured RNA substrates. Ligation of the linear ASL-4U substrates (l) to the circular form (c) was monitored by urea PAGE. The bracket marks formed concatemers (concat.).

B, EMSA with WT *Ct*Trl1-LIG (0-10 µM) and ASL-4U or ASL-4U$^{-6}$ (1 µM). Increasing amounts of proteins were incubated with the RNA and analysed by native PAGE. Formation of the LIG•RNA complex was assessed by band shift.

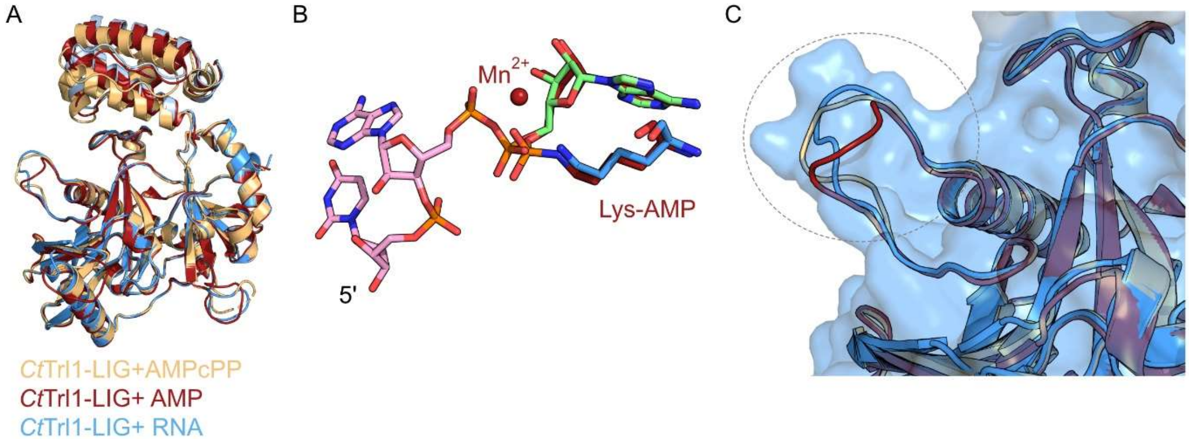


**Extended Data Fig. 3: Structural alignment of *Ct*Trl1-LIG•RNA with other *Ct*Trl1-LIG structures.**

A, Superposition of *Ct*Trl1-LIG domain crystal structures in complex with AMPcPP (orange, PDB ID: 6N67), AMP (red, PDB ID: 6N0V) and with RNA (blue, PDB ID: 8RBJ). The structures were superimposed to the β-sheets of the adenylyltransferase domain (β5, β6, β8, β9, β14 and β15).

B, Zoom into the position of the active site lysine K148N relative to AMP moiety and activated RNA end of the *Ct*Trl1-LIG•AMP structure (red) and the *Ct*Trl1-LIG•RNA structure.

C, Zoom into the thumb-like loop area (S168-S181) in the superposition of *Ct*Trl1-LIG structures. The surface of the *Ct*Trl1-LIG•RNA is presented transparently. The dashed circle marks the thumb-like protrusion. The color code is the same as in A.

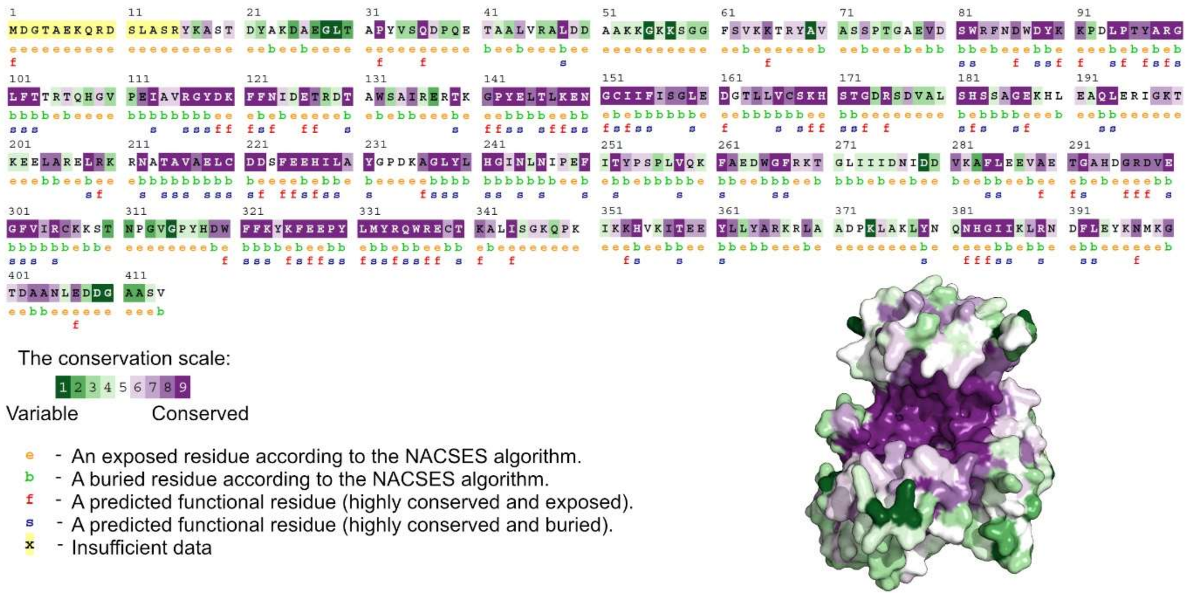


**Extended Data Fig. 4: Conservation analysis of the Trl1-LIG domain.**
The degree of amino acid conservation was determined using the ConSurf server[1,2]. The *Ct*Trl1 sequence was used as input to calculate conservation of the residues in the LIG domain among Trl1-type ligases (with all parameters set to default). The conservation scale indicates the degree of conservation from most variable (green; 1) to most conserved (purple; 9). Additional residue information is given in the legend. The Multiple sequence alignment as well as the *Ct*Trl1-LIG structure are shown in the conservation scale colors, respectively.

A

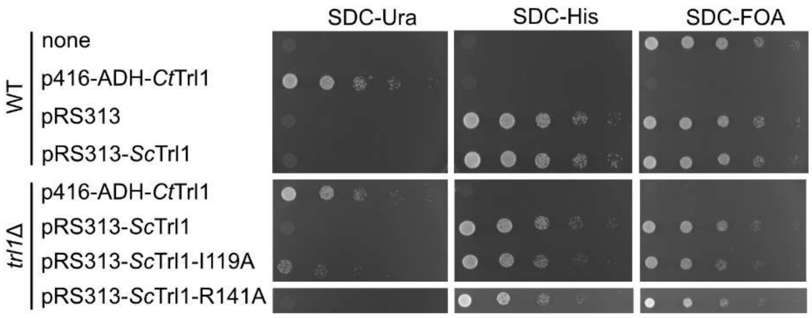


B

| RNA strand | *Ct*Trl1 | *Sc*Trl1 | Mutation to alanine |
|---|---|---|---|
| 5' exon | K64 | - | - |
| | Y68 | - | - |
| | R83 | - | - |
| | I153 | I119 | Viable |
| | S168 | S134 | Lethal* |
| | K169 | K135 | Lethal* |
| | H170 | H136 | Lethal* |
| | S171 | S137 | Lethal* |
| | R175 | R141 | Viable |
| | H182 | H148 | Lethal* |
| adenylylated 3' exon | R99 | R66 | Lethal* |
| | K169 | K135 | Lethal* |
| | K323 | K284 | Lethal* |
| | K325 | K286 | Lethal* |
| 3' exon in duplex | N382 | - | - |
| 5' exon in duplex | No interactions found. | | |

*Wang and Shuman, 2005

**Extended Data Fig. 5: Impact of RNA-coordinating resides on yeast cell growth.**
A, Yeast growth assay of the indicated strains to test cell survival of controls and after introducing Trl1 mutations. Dilution series (5-fold) of yeast cells were spotted on the indicated selective solid media.
B, Overview of RNA binding amino acids in *Ct*Trl1 (based on PDB ID: 8RBJ) as well as the corresponding residues in yeast (*Sc*Trl1). The cell viability of the respective alanine variants in *S. cerevisiae* is indicated based on this study or in[3].

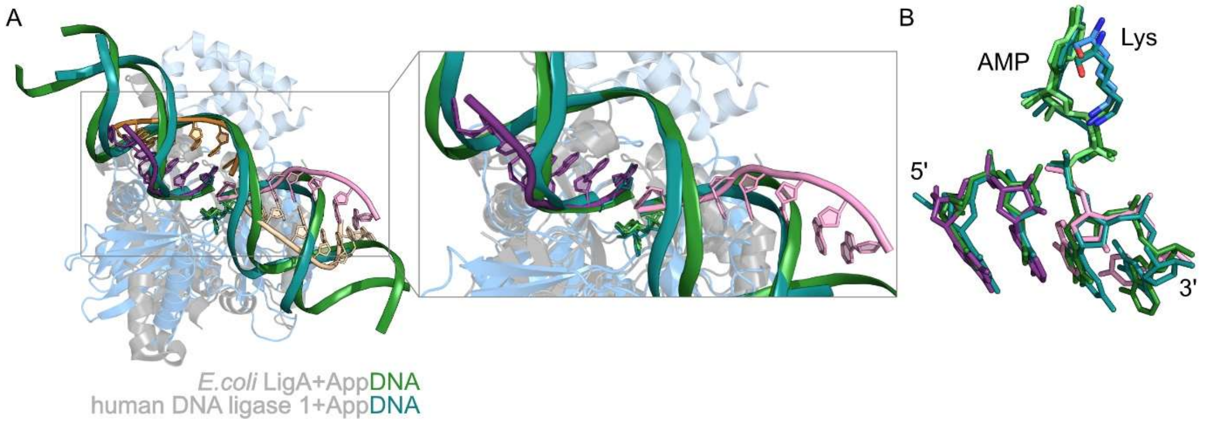


**Extended Data Fig. 6: Superposition of *Ct*Trl1-LIG•RNA with *DNA ligase*•AppDNA structures reveals conserved modes of RNA binding.**

A, Superposition of our *Ct*Trl1-LIG•RNA structure with the crystal structures of *E. coli* LigA•AppDNA (PDB ID: 2OWO, grey protein and dark green DNA strands) and human DNA ligase 1•AppDNA (PDB ID: 1X9N, grey protein and dark cyan DNA strands). The structures were superposed to the β-sheets of the adenylyltransferase domain. The DNA strands are depicted as cartoons.

B, Zoom into the activated 5′ end of the 3′ exon and the 3′ end of the 5′ exon of the nucleic acids in the structures from A reveals a conserved coordination principle among adenylyltransferases. The active site lysine residues (K148 of *Ct*Trl1-LIG, K115 of *E. coli* LigA and K568 of the human DNA ligase 1) as well as the first two nucleotides of both strands are depicted as sticks.

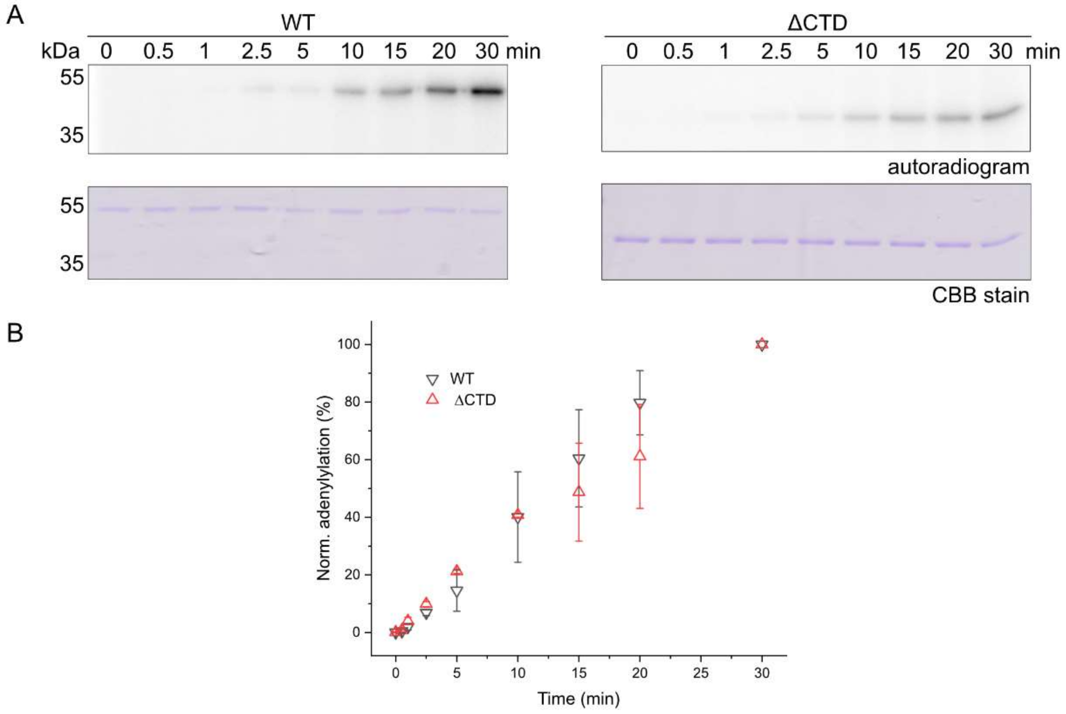


**Extended Data Fig. 7: Time-dependent active site adenylylation of *Ct*Trl1-LIG WT and ΔCTD.**
A, Active site adenylylation of WT *Ct*Trl1-LIG and the ΔCTD variant was monitored by incorporation of $\alpha P^{32}$-labeled ATP. The time course of the adenylylation reaction was analyzed by autoradiogram (upper panel). Even protein loading was confirmed by SDS-PAGE and Coomassie Brilliant Blue (CBB) staining (lower panel).
B, Quantification of the experiment in A performed as triplicates. Data are depicted as mean including standard deviation.

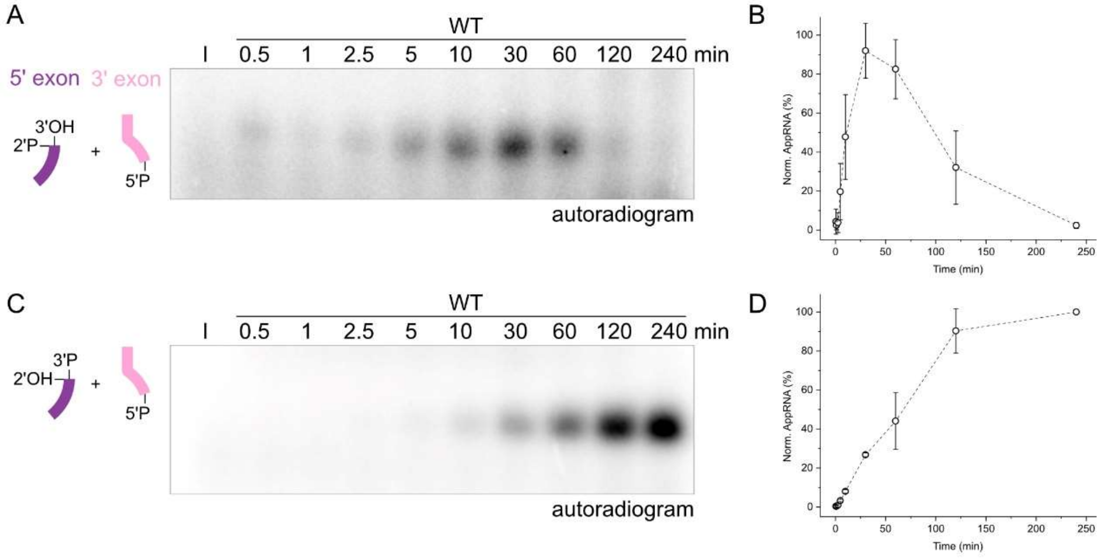


**Extended Data Fig. 8: Time-dependent formation of the AppRNA intermediate.**

A, Time course of RNA (2′-P and 3′-OH on the 5′ exon end and 5′-P on the 3′ exon end) adenylylation by *Ct*Trl1-LIG WT. The autoradiogram shows formation of the adenylylated AMP-ASL-4U intermediate via $\alpha P^{32}$-AMP incorporation.

B, Densitometric quantification of the experiment in A. The graph reveals a bell-shaped curve for the formation of the AppRNA intermediate over time and its fading through the ligation reaction.

C, Time course of RNA (2′-OH and 3′-P on the 5′ exon end and 5′-P on the 3′ exon end) adenylylation by *Ct*Trl1-LIG WT. The autoradiogram shows formation of the adenylylated AMP-ASL-4U intermediate via $\alpha P^{32}$-AMP incorporation.

D, Densitometric quantification of the experiment in C. The graph reveals a saturation curve for the formation of the AppRNA intermediate over time.

**Extended Data Table 1: Oligonucleotides for site-directed mutagenesis of pET15b-ctTrl1-LIG-WT**

| Name | Sequence 5′→3′ | Source |
|---|---|---|
| ctTRL1-K148N_sdm | GTACGAGCTCACACTCAACGAGAACGGCTGCATC | Sigma |
| ctTRL1-E328Stop_sdm | CAAGTACAAGTTCGAGTAACCTTACCTGATG | Sigma |
| ctTrl1-LIG-R334A_sdm-f | CCTGATGTACGCGCAGTGGCGTG | Sigma |
| ctTrl1-LIG-R334A_sdm-r | TAAGGTTCCTCGAACTTG | Sigma |
| ctTrl1-LIG-R337A_sdm-f | CCGGCAGTGGGCTGAATGTACC | Sigma |
| ctTrl1-LIG-R337A_sdm-r | TACATCAGGTAAGGTTCC | Sigma |
| ctTrl1-LIG-R334.337A_f | CGCGCAGTGGGCTGAATGTACC | Sigma |

**Extended Data Table 2: RNA substrates**

| Name | Sequence 5′→3′ | Source |
|---|---|---|
| sctRNA$^{Phe}$-3′exon-7nt | [Phos]AUCUGGA | Sigma |
| ASL-4U | [Phos]AUCUGGAUUUUGCCAGACUGAAG[Phos] | Sigma |
| ASL-4U$^{Inv}$ | [Phos]GAAGUCAGACCGUUUUAGGUCUA[Phos] | Sigma |
| ASL-4U$^{AU}$ | [Phos]AUAUAUAUUUUGAUAUACUGAAG[Phos] | Sigma |
| ASL-4U$^{AG}$ | [Phos]AAGAGAGAGAGAGGAGAGAGAAA[Phos] | Sigma |
| ASL-4U$^{-2nt}$ | [Phos]AUCUGGAUUUUGCCAGACUGA[Phos] | Sigma |
| ASL-4U$^{-4nt}$ | [Phos]AUCUGGAUUUUGCCAGACU[Phos] | Sigma |
| ASL-4U$^{-6nt}$ | [Phos]AUCUGGAUUUUGCCAGA[Phos] | Sigma |
| ASL-4U$^{-3'P}$ | [Phos]AUCUGGAUUUUGCCAGACUGAAG | Sigma |
| HAC1-4U | [Phos]AAGCGCGAUUUUGCGCGUAAUCCAG[Phos] | Sigma |

**Extended Data Table 3: Oligonucleotides for yeast expression plasmid construction**

| Name | Sequence 5′→3′ | Source |
|---|---|---|
| Fragment1_f | CTAGAACTAGTGGATCCCCCAAAGGTTCAAGGAATGCCATGC | Sigma |
| Fragment1_r | GGCTAGGCATCGCTTCTTCGTATGAATACTTTTATGATCACTAAAG | Sigma |
| Fragment2_f | CGAAGAAGCGATGCCTAGCCC | Sigma |
| Fragment2_r | AGACCGCGGTCTAGGCGGCCGG | Sigma |
| Fragment3_f | GCCGCCTAGACCGCGGTCTGGCT | Sigma |
| Fragment3_r | TATCGATAAGCTTGATATCGTAGAACGAGAATTTCGACCGAAGGTG | Sigma |

**Extended Data Table 4: Oligonucleotides for site-directed mutagenesis of pRS313-*Sc*Trl1**

| Name | Sequence 5′→3′ | Source |
|---|---|---|
| pRS313-NotI-Trl1_fwd | GGTGGCGGCCGCTCTAGAACT | Sigma |
| Trl1-SalI-pRS313-rev | TCGAGGTCGACGGTATCGATAAGC | Sigma |
| scTrl1-I119A_sdm-f | CAATGGTTGTgcCATTTTTATATCTGG | Sigma |
| scTrl1-I119A_sdm-r | GCTTTTATAGTGACATCG | Sigma |
| scTrl1-R141A_sdm-f | TACTGGACCCgcAGCAGACGTAG | Sigma |
| scTrl1-R141A_sdm-r | GAATGCTTCGAACAAAC | Sigma |

**Extended Data Table 5: Yeast expression plasmids**

| Name | Auxotrophy marker | Source |
|---|---|---|
| p416-ADH-ctTrl1 | Ura | [4] |
| pRS313 | His | [5] |
| pRS313-scTrl1-3xHA | His | This study |
| pRS313-scTrl1-I119A-3xHA | His | This study |
| pRS313-scTrl1-R141A-3xHA | His | This study |

**Extended Data Table 6: Yeast strains**

| Yeast strain | Auxotrophy marker | Source |
|---|---|---|
| WT, W303<br>*(MATα leu2-3,112 TRP1 can1-100 ura3-1 ADE2 his3-11,15)* | - | [4] |
| WT + p416-ADH-ctTrl1 | Ura | This study |
| WT + p416-ADH-ctTrl1 + pRS313-scTrl1-3xHA | Ura, His | This study |
| WT + pRS313 | His | This study |
| WT + pRS313-scTrl1-3xHA | His | This study |
| Δtrl1 + p416-ADH-ctTrl1 | Ura | [4] |
| Δtrl1 + p416-ADH-ctTrl1 + pRS313-scTrl1-3xHA | Ura, His | This study |
| Δtrl1 + pRS313-scTrl1-3xHA | His | This study |
| Δtrl1 + pRS313-scTrl1-I119A-3xHA | His | This study |
| Δtrl1 + pRS313-scTrl1-R141A-3xHA | His | This study |
